# Characterizing a hallmark of glymphatic insufficiency: Wasteosomes accumulate in periventricular white matter hyperintensities and exhibit complex relationships with mixed pathology, sclerotic index and perivascular space

**DOI:** 10.1101/2025.07.25.666856

**Authors:** Nikita Ollen-Bittle, Isabella Boesgaard, Austyn Roseborough, Meaghan Frank, Qi Zhang, Stephen H. Pasternak, Robert Hammond, Shawn N. Whitehead

**Affiliations:** Department of Anatomy and Cell Biology, Schulich School of Medicine and Dentistry, Western University, London, Ontario, N6A 5C1; Robarts Research Institute, Schulich School of Medicine and Dentistry, London, ON, Canada; Department of Clinical Neurological Sciences, Schulich School of Medicine and Dentistry, Western University, London, Ontario, N6A 5C1; Department of Pathology and Laboratory Medicine, London Health Sciences Centre, London, ON, Canada

**Keywords:** Wasteosomes, corpora amylacea, white matter hyperintensities, Alzheimer’s disease, cerebrovascular disease, glymphatic system

## Abstract

The glymphatic system is a recently elucidated waste clearance system in the brain, thought to be critical for the maintenance of homeostasis. Corpora amylacea or “wasteosomes”, are discontinuous lipid labyrinth structures that are polyglucosan rich, retain cellular waste and are thought to be of astrocytic origin. Wasteosomes have been proposed as a hallmark of glymphatic insufficiency predominantly due to: 1) their spatial localization around glymphatic drainage points including periventricular (PV) regions, perivascular spaces (PVS), and sub-pial regions; and, 2) their correlation with aging, vascular disorders, neurodegenerative diseases, and conditions that impair sleep. White matter hyperintensities (WMHs) are diffuse hyperintense areas seen on T2-weighted or fluid-attenuated inversion recovery (FLAIR) magnetic resonance imaging (MRI) scans that represent damage to white matter. PV WMHs and are known predictors of mild cognitive impairment, stroke, dementia and death. The relationship between wasteosome accumulation, PV WMHs, vascular pathology and PVS is currently unknown. For the first time, in a mixed diagnostic cohort of pathologically diagnosed: Alzheimer’s disease (AD), cerebrovascular disease (CVD), mixed AD/CVD, and control tissue with no pathological diagnosis, we connected the histopathological wasteosome profile in periventricular brain sections in relation to 7T FLAIR-MRI confirmed PV WMHs, vascular stenosis and PVS. Our results reveal wasteosomes accumulate in PV WMHs, are increased in proximity to large PV venules, and exhibit complex relationships with WMH severity, mixed pathology, sclerotic index and PVS. These findings suggest wasteosomes may serve as histological markers of impaired glymphatic drainage and provide new insights into the pathophysiology underlying white matter injury.

## Introduction

Corpora amylacea, hereafter referred to as wasteosomes, are discontinuous lipid labyrinth structures that are polyglucosan rich and retain cellular waste. Wasteosomes range in size from 10-50 μm in diameter and are found throughout the body including in the brain^1^. Within the brain wasteosomes have been shown contain elements of neuronal, astrocytic, oligodendrocytic, hematological and even infectious origins; however, the vast majority of the scientific literature suggests they are primarily produced by astrocytes^2,3^. Wasteosomes are thought to serve as protective waste containers, shielding the brain’s functional architecture from a variety of harmful lipid, protein and microbial biomolecules that may be non-degradable otherwise^4,5^. While their exact composition has been a topic of debate, consensus notes wasteosomes are predominantly composed of polymerized hexoses which account for approximately 88% of their dry weight^6,7^, and contain neoepitopes recognized by natural antibodies for the immunoglobulin M (IgM) isotope^3^. Wasteosomes stain a bright magenta with periodic acid schiff (PAS) staining and can also be detected as a blue/green colour with alcian blue positivity in Movat’s pentachrome staining. Some wasteosomes have been found to exhibit a central core as well as a peripheral region containing glycogen synthase, a requisite enzyme for polyglucosan formation, ubiquitin and protein p62 which are known to be associated with waste clearance^3^. This has led to the hypothesis that the central core may correspond to a region of ubiquinated waste build-up, while the peripheral region may be an active zone whereby p62 collects waste to be packaged by glycogen synthase^3^. Moreover, recent work has demonstrated the composition of wasteosomes may vary between disease states. A 2023 study by Riba and colleagues found tau protein to be present in some wasteosomes in Alzheimer’s disease (AD) patients, but not in non-AD patients^8^. The composition of wasteosomes is only beginning to be elucidated, but emerging evidence widely supports wasteosomes as a key component of waste removal in the brain.

In addition to wasteosome composition, the neuroanatomical localization of wasteosomes further infers their role as waste bodies to be cleared from the brain parenchyma. Wasteosomes are predominantly found to concentrate at perivascular, subpial, and peri-ventricular regions of the brain. In 1969, Sakai and colleagues mapped the distribution of wasteosomes in the cerebrum of four brains from 70-year-old donors and found wasteosomes predominantly aggregated in brain regions close to cerebrospinal fluid (CSF) including the walls of the ventricles and deep regions of cerebral sulci^7^. The wasteosome map developed by these authors is now known to closely correlate with neuroanatomical regions critically implicated in glymphatic drainage^9^. The glymphatic system in the brain is thought to serve an analogous function to the lymphatic drainage system throughout other organs in the body. The glymphatic system functions to drain interstitial fluid (ISF) and remove metabolic and neurotoxic waste. Flow through the glymphatic system is in part driven through the pulsatile nature of arteriole vessels and seemingly follows a circadian rhythm increasing during sleep^10^. The glymphatic system involves the flow of CSF from the subarachnoid space in the periarterial spaces where it is propelled along and astrocytic endfeet via aquaporin-4 (AQP4) to facilitate mixing with brain ISF, such that it then flows, largely following white matter tracts, to exit the brain via the ventricles, perivenous spaces, and even spaces surrounding cranial nerves^9^. Intriguingly, just as aging, sleep disorders and cerebral vascular disease are known to directly impact the glymphatic system, the relative abundance of wasteosomes in the brain has been shown to increase with age^3^, certain vascular disorders, neurodegenerative diseases^8^ and conditions that impair sleep such as sleep apnea^11^. The correlation between wasteosome accumulation, vascular dysregulation, age, and disordered sleep has recently led to wasteosomes being proposed as a hallmark of glymphatic insufficiency^9^. Collectively, wasteosomes represent a poorly understood entity that may directly relate to cerebral vascular pathology and impaired glymphatic clearance.

White matter hyperintensities (WMHs) are diffuse hyperintense areas seen on T2-weighted or fluid-attenuated inversion recovery (FLAIR) magnetic resonance imaging (MRI) scans representing damage to white matter^12^. WMHs are synonymous with leukoariosis reported on computed tomography (CT) scans^13^ and are extremely common findings with advancing age, occurring in the majority of individuals over 60 years of age^14^. Clinically, WMHs are often thought of as a signal of cerebral small vessel disease^15^ and concurrent periventricular infarcts (PVI) in some WMHs would suggest the same^16^. Importantly, greater WMH volume is a known predictor of mild cognitive impairment, stroke, dementia and death^17^ and WMH pathology has been closely associated with a lower threshold for clinically overt dementia^18^. White matter homeostasis is thought to rely heavily on functionality of the brain’s glymphatic system and ISF flow to remove metabolic byproducts from the extensive network of myelinated axons^19^. Moreover, as previously noted, ISF flow has been shown to follow white matter tracts towards perivenous spaces^9,20^. Despite the intriguing relationship between WMHs, vascular dysregulation and glymphatic flow, the relationship between wasteosome accumulation and WMHs, vascular stenosis and perivascular space (PVS) is currently unknown. We hypothesized wasteosome accumulation would colocalize with WMHs and correlate with pathological diagnosis and indicators of cerebrovascular pathology including PVS enlargement and vascular stenosis. To address this question, in a mixed specimen cohort of pathologically diagnosed: AD, cerebrovascular disease (CVD), mixed AD/CVD, and control tissue with no pathological diagnosis, we assessed the wasteosome profile in periventricular (PV) brain sections in relation to 7T FLAIR MRI confirmed WMHs, vascular stenosis and PVS.

## Methods

### Specimen selection and tissue handling

All specimens analyzed in this study were derived from a cohort previously characterized by Roseborough et al. (**Table 1**)^16^. In brief, 20 specimens were selected and grouped based on neuropathological diagnoses including: five pathologically normal, five AD, five CVD and five mixed AD/CVD. Following MRI, one histological tissue block from both the left and right hemisphere were sampled by neuropathologists to include PV white matter, subcortical (SC) white matter and cortical tissue. Tissue blocks were placed into histological cassettes and stored in 10% formalin. Of note, one tissue block previously utilized in Roseborough et al. 2021 was unable to be utilized for this publication. All other specimens were included.

**Table 1.** Demographic and pathological details. All demographic details available for donor specimens included in this study. Specimens originally characterized and table originally published in Roseborough et al. 2021^16^.

| Specimen | Age | Sex | Pathology Diagnosis | Fixation (mo.) |
| --- | --- | --- | --- | --- |
| 1 | 75 | M | Mild AD (Braak stage 1/2) | 158 |
| 2 | 71 | M | Advanced AD (Braak stage 5/6), NFT, CAA, superficial siderosis | 151 |
| 3 | 80 | F | Advanced AD (Braak stage 5/6) | 155 |
| 4 | 68 | F | Mild AD (Braak stage 1/2) | 155 |
| 5 | 83 | M | Advanced AD (Braak stage 5/6) | 160 |
| 6 | 73 | F | Advanced AD (Braak stage 5/6), CVD, acute infarct, vasculopathy | 161 |
| 7 | 91 | F | Advanced AD (Braak 5/6), cerebrovascular disease, atherosclerosis | 155 |
| 8 | 84 | M | Old infarct cerebellum, mild AD (Braak stage 1/2) | 56 |
| 9 | 80 | F | Advanced AD (Braak 5/6), CVD | 154 |
| 10 | 83 | F | Advanced AD (Braak stage 5/6), diffuse<br>Lewy body, CVD, CAA | 155 |
| 11 | 78 | M | Acute microinfarcts | 151 |
| 12 | 75 | M | Cerebrovascular disease & old bleed | 154 |
| 13 | 78 | F | Multiple infarcts | 148 |
| 14 | 47 | M | Multiple infarcts, CVD, arteriosclerosis | 157 |
| 15 | 85 | M | CVD, Crookes hyalinization, acute HIE<br>CA1 | 162 |
| 16 | 65 | F | No significant findings | 151 |
| 17 | 52 | F | No significant findings | 158 |
| 18 | 69 | M | No significant findings | 164 |
| 19 | 70 | M | No significant findings | 159 |
| 20 | 52 | F | No significant findings | 161 |

### Magnetic resonance imaging

MRI scans were conducted using a 7T Siemens Scanner at Western’s Centre for Functional and Metabolic Mapping according to a previously described protocol^21^. In brief, frontal coronal sections from each fixed brain were imaged in Galden HT-270 perfluorinated fluid in a custom-stacking device and T1 (T1-weighted, MP2RAGE sequence with extended T1 time to improve contrast) and T2-FLAIR (T2 SPACE sequence) images were acquired.

### Evaluation of white matter hyperintensities

Details on the assessment of WMHs in these specimens has previously been published^16^. Briefly, all assessment of WMHs was done blinded to neuropathological diagnosis and histological data. PV WMHs were assessed on FLAIR sequence utilizing Fazekas criteria^22^ and rated as either none, mild (extending outwards <1/3 of the SC white matter), moderate (>1/3 but < 2/3 of the SC white matter) OR as having a PVI based on T1-weighted imaging.

### Histology and immunohistochemistry

All immunohistochemistry (IHC) and histological staining was done by the Department of Pathology and Laboratory Medicine, University Hospital, London, Ontario, Canada. Tissue was paraffin embedded and sectioned at 8 μm for staining and 4 μm for IHC. A detailed protocol for tissue staining has previously been published^16^. For the purposes of this study, Movat’s pentachrome staining was utilized for the visualization of vascular architecture and wasteosome accumulation. Smooth muscle actin IHC was utilized to discern concentric rings of smooth muscle cells in the tunica media of arterioles.

### Stenosis, perivascular space and average wasteosome number per vessel

Slides were scanned with an Aperio Whole-Slide Scanner by Pathology Core Facilities at Western University. Aperio ImageScope Software was then utilized to measure blood vessels and count wasteosomes. Calculation of sclerotic index (SI) and PVS were performed as previously described, with all measurements taken from the maximum vessel diameter^23^. Vessels were grouped based on arteriole or venule classification, anatomical location and size. Regarding anatomical location vessels were grouped as either: 1) PV vessels, located within 0.5 cm from the ependymal lining of the ventricle; or 2) SC vessels, any vessels outside of the 0.5 cm boundary. Vessels were further subdivided based on size with medium (50 μm < IntD < 200 μm) and large (>200 μm) vessels measured. Wasteosomes were counted within a 200 μm radius of their corresponding vessels. In the case where vessels were close in proximity and the 200 μm radius overlapped, wasteosomes were counted with whichever vessel they were most proximal to. As in Roseborough et al. 2021^16^, three vessels were measured for each size group whenever possible and average SI, PVS and wasteosome count per vessel were used for analysis.

### Assessment of wasteosome accumulation in WMHs

Tissue blocks corresponding to moderate and PVI groups were combined for assessment of wasteosome accumulation in WMHs. Only specimens within these groups with both a PV WMH and an area of PV normal appearing white matter (NAWM) were utilized. Specimens with only SC NAWM were not included in this analysis. Regions of interest (ROIs) of approximately the same sizes were annotated around NAWM and WMHs utilizing Aperio ImageScope Software based on FLAIR MRI sequences and wasteosomes were counted within these ROIs.

### Statistics

WMH vs NAWM wasteosome accumulation: Normality was assessed with a Shapiro-Wilk normality test and determined to be non-parametric. Wasteosome accumulation was then assessed utilizing a Wilcoxon matched-pairs signed rank test.

Other analyses, greater than two groups: Normality was assessed with a Shapiro-Wilk normality test. Parametric data was analyzed using an ANOVA test with Tukey post-hoc, while non-parametric data was analyzed with a Kruskal-Wallis test with Dunn’s post-hoc.

Correlation analysis: Normality was first assessed with a Shapiro-Wilk normality test. Parametric data was analyzed using Pearson correlation coefficients and simple linear regression while non-parametric data was analyzed with a Spearman correlation.

Significance was determined when p < 0.05. All statistics were performed using GraphPad Prism Software Version 10.5.0.

### Graphical Abstract

Created with BioRender.com

## Results

To characterize wasteosome localization in PV white matter, we performed systematic histological analysis. This revealed a distinct pattern of wasteosome accumulation, with predominant clustering around blood vessels (**Figure 1A**) and increased density in PV compared to SC regions. First, we assessed wasteosome accumulation in MRI confirmed WMHs in specimens either scored as moderate WMH or as having a PVI, compared to adjacent NAWM. To mitigate the subjective observation of substantially more wasteosomes in the PV region, only specimens with both PV WMHs and NAWM were included in this analysis. **Figure 1B** depicts an MRI confirmed WMH region as well as ROIs within the WMH and within the adjacent NAWM. No significant differences were found when comparing wasteosome accumulation between these groups (**Figure 1C).** The results between these groups were then pooled and paired analysis between NAWM and WMH ROIs within the same specimen revealed significantly increased wasteosome accumulation in WMHs (**Figure 1D**). We then sought to characterize the subclass of blood vessels that wasteosomes tend to accumulate around in both PV and SC regions. First assessing large vessels, results demonstrated significantly more wasteosomes accumulated around large PV venules compared to PV arterioles and SC arterioles and venules (**Figure 1E**, p < 0.0001). Contrastingly, in medium sized vessels, no significant difference in wasteosome accumulation was observed around PV venules and arterioles. However, more wasteosomes were found to accumulate around medium sized PV venules compared to medium sized SC venules (p = 0.0007) and arterioles (p = 0.0076), (**Figure 1F**). To interrogate if the differing results between large and medium sized vessels could be explained by the size of the PVSs surrounding the vessels, we then normalized the average number of wasteosomes per vessel by the average PVS. In large vessels, results still revealed significantly more wasteosomes accumulated around large PV venules compared to PV arterioles and SC arterioles and venules (p < 0.0001), and also revealed significantly more (p = 0.0470) wasteosomes accumulated around large SC venules compared to SC arterioles (**Figure 1G**). In the medium sized vessels significantly more wasteosomes were found to accumulate around PV arterioles (p = 0.0338) and venules (p = 0.0024) compared to their SC counterparts, however, no differences were observed between PV arterioles and venules or SC arterioles and venules (**Figure 1H**).

**Figure 1.**
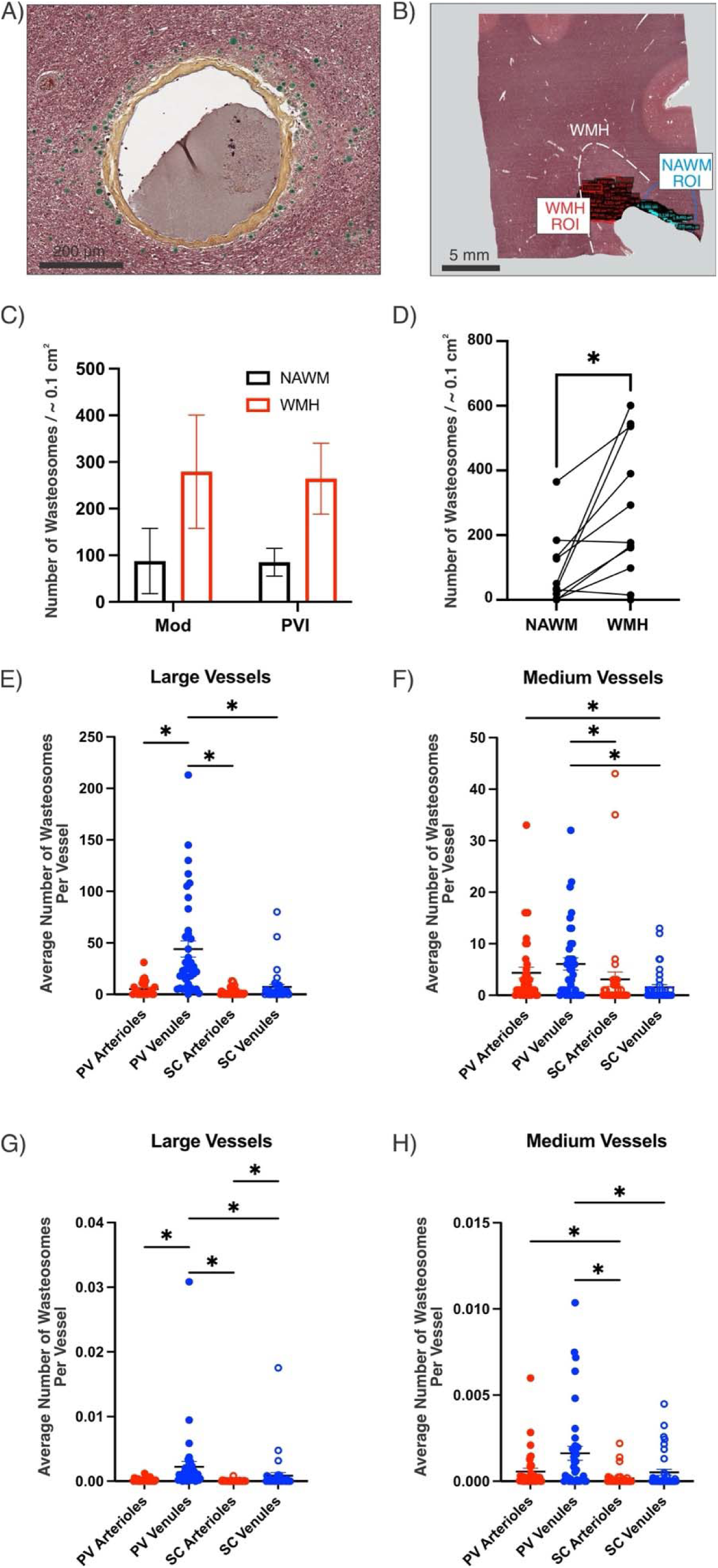
Wasteosomes accumulate in white matter hyperintensities and localize around large periventricular venules. A) Example of wasteosomes accumulating around a partially stenosed periventricular (PV) blood vessel. B) Movat’s pentachrome stained tissue section with regions of interest (ROIs) drawn out in both a magnetic resonance imaging (MRI) confirmed white matter hyperintensity (WMH) and region of normal appearing white matter (NAWM). C) Wasteosomes were counted in tissue sections with WMHs classified as either moderate (Mod), or as having a periventricular infarct (PVI). No significant difference was found between groups. D) The number of wasteosomes in WMHs and NAWM regions in sections classified as Mod and PVI were pooled and paired analysis revealed significantly greater wasteosome accumulation in WMHs relative to adjacent NAWM (p = 0.0127). E) Significantly more wasteosomes accumulate around large PV venules compared to PV arterioles and SC arterioles and venules (p < 0.0001, n PV Arterioles = 26, n PV Venules = 38, n SC Arterioles = 38, n SC Venules = 38). F) There is no difference in wasteosome accumulation around medium sized PV venules and arterioles; however more wasteosomes accumulate around medium sized PV venules compared to medium sized SC venules (p = 0.0007) and arterioles (p = 0.0076), (PV Arterioles = 38, n PV Venules = 38, n SC Arterioles = 38, n SC Venules = 38). G) Average number of wasteosomes per vessel were then normalized by PVS. In Large vessels significantly more wasteosomes accumulated around large PV venules compared to PV arterioles and SC arterioles and venules (p < 0.0001), and significantly more (p = 0.0470) wasteosomes accumulated around large SC venules compared to SC arterioles. H) In the medium sized vessels significantly more wasteosomes accumulated around PV arterioles (p = 0.0338) and venules (p = 0.0024) compared to their SC counterparts, however, no differences were observed between PV arterioles and venules or SC arterioles and venules.

To determine the impact of disease severity and pathological classification on perivascular wasteosome distribution, we analyzed accumulation patterns across different vessel subtypes stratified by WMH scoring and diagnostic categories. Wasteosome accumulation around both medium and large blood vessels was found to be independent of WMH scoring (**Figure 2A-H)**. Pathological diagnosis was also largely found to have no influence on wasteosome accumulation around blood vessels, with the exception of medium sized SC arterioles (**Figure 2I-P**). A pathological diagnosis of mixed AD/CVD pathology was found to significantly increase (p = 0.0229) wasteosome accumulation around medium sized SC arterioles relative to specimens with a pathological diagnosis of CVD alone (**Figure 2O**).

**Figure 2.**
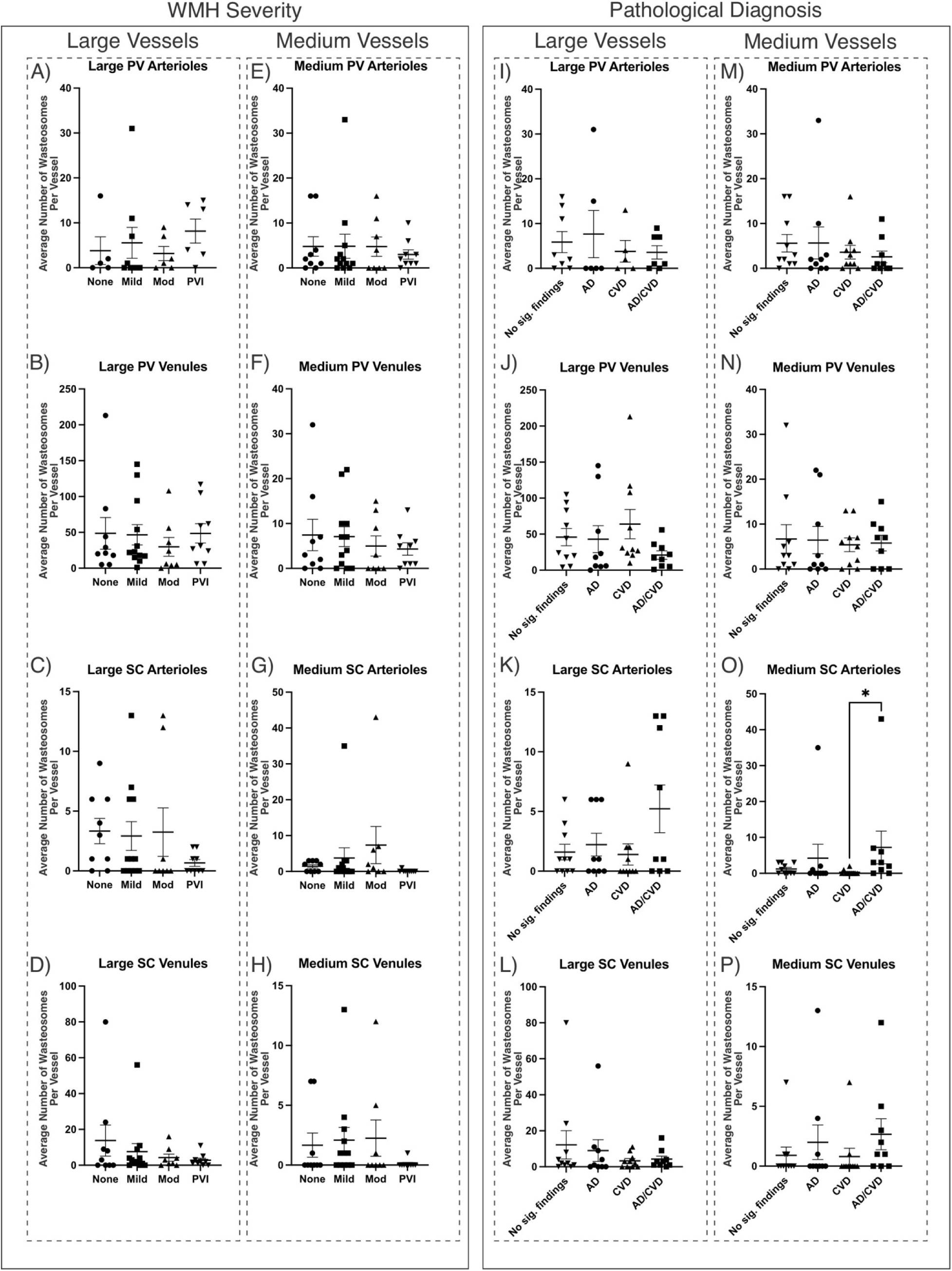
Wasteosome accumulation around blood vessels is independent of WMH severity but may be influenced by comorbid pathology. A-D) No significant differences were found in the average number of wasteosomes per vessel in large periventricular (PV) and subcortical (SC) vessels between severities of white matter hyperintensities (WMHs). E-H) No significant differences were found in the average number of wasteosomes per vessel in medium PV and SC vessels between severities of WMHs. I-L) Pathological diagnosis did not affect the average number of wasteosomes per vessels in large PV and SC vessels. M-N) Pathological diagnosis did not affect the average number of wasteosomes per vessels in medium PV vessels; however, O) a pathological diagnosis of AD/CVD significantly increases (p = 0.0229) wasteosome accumulation around medium sized SC arterioles relative to those in specimens with a pathological diagnosis of CVD alone. P) Although a similar trend was observed in medium sized SC venules, no significant difference was found.

Next, we examined the relationship between SI, PVS and wasteosome accumulation around blood vessels. Intriguingly, an inverse relationship between wasteosome accumulation and SI was observed in distinct medium sized vessels. Negative correlations between wasteosome accumulation and SI were found in: 1) medium sized PV venules in specimens with no WMHs (r = -0.6946, p = 0.0448); 2) medium sized PV arterioles in specimens with no significant pathological findings (r = -0.8441, p = 0.0448); and, 3) in medium sized SC venules within specimens with a pathological diagnosis of AD (r = -0.7327, p = 0.0357). Analysis of other vessel types, diagnostic groups and WMH scoring groups revealed no other significant findings. **Figure 3** showcases all significant findings.

**Figure 3.**
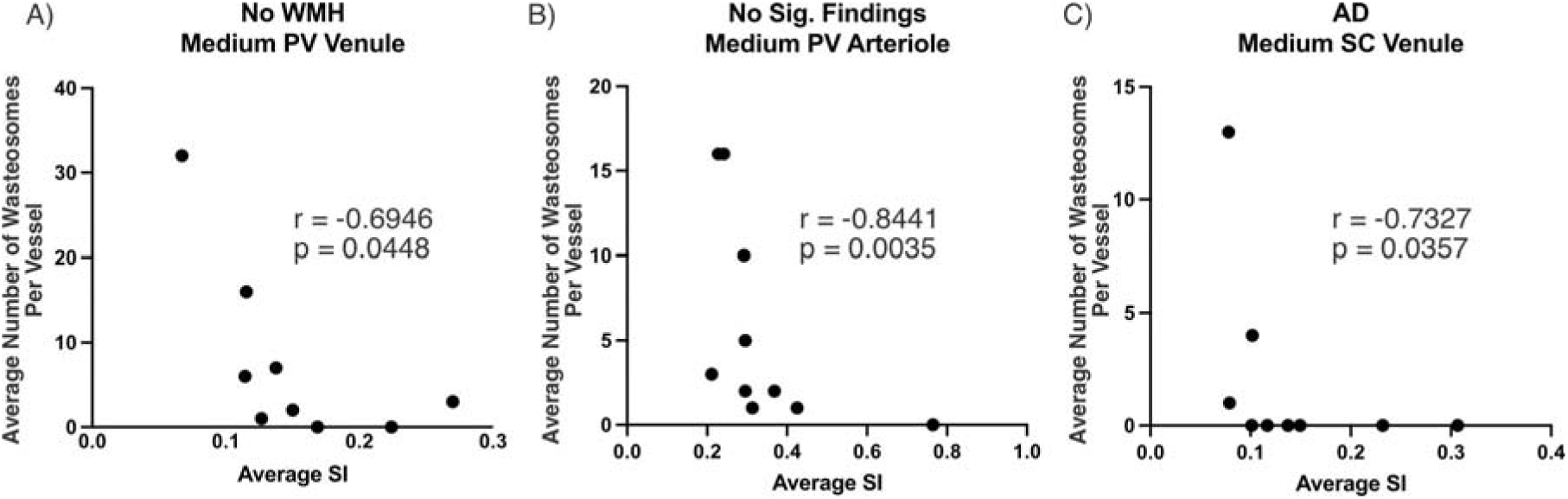
Evidence of an inverse relationship between wasteosome accumulation and the sclerotic index of medium sized vessels. A) In specimens with no magnetic resonance imaging (MRI) confirmed white matter hyperintensities (WMHs), the average wasteosome accumulation around medium sized periventricular (PV) venules is negatively correlated with the average sclerotic index (SI) of those vessels. B) This relationship was further found in medium sized PV arterioles in specimens grouped based on no significant pathological findings, and C) in medium sized subcortical (SC) venules within specimens with a pathological diagnosis of AD.

To elucidate the associations between vascular structural parameters and PV waste accumulation, we analyzed the relationships between SI, PVS, and wasteosome density around different blood vessel subtypes. Despite an insignificant correlation in specimens with no WMHs around large PV venules (**Figure 4A**), and a significant positive correlation in specimens with mild WMHs around medium subcortical (SC) arterioles (r = 0.7195, p = 0.0114) (**Figure 4B**), a significant negative correlation was found in specimens with moderate WMHs between PVS and wasteosome accumulation around large PV venules (r = -0.7619, p = 0.0368) (**Figure 4C**). A negative relationship was also observed in specimens with AD/CVD pathological diagnoses between wasteosome accumulation and PVS of large PV venules (r = -0.6974, R^2^ = 0.4864, p = 0.0368) (**Figure 4D**). However, positive relationships between PVS and wasteosome accumulation in specimens with AD/CVD were found in medium PV venules (r = 0.8421, R2 = 0.7091, p = 0.0044) (**Figure 4E**), and large SC arterioles (r = 0.9063, p = 0.0020) (**Figure 4F**). Analysis of other vessel types, diagnostic groups and WMH scoring groups revealed no other significant findings.

**Figure 4.**
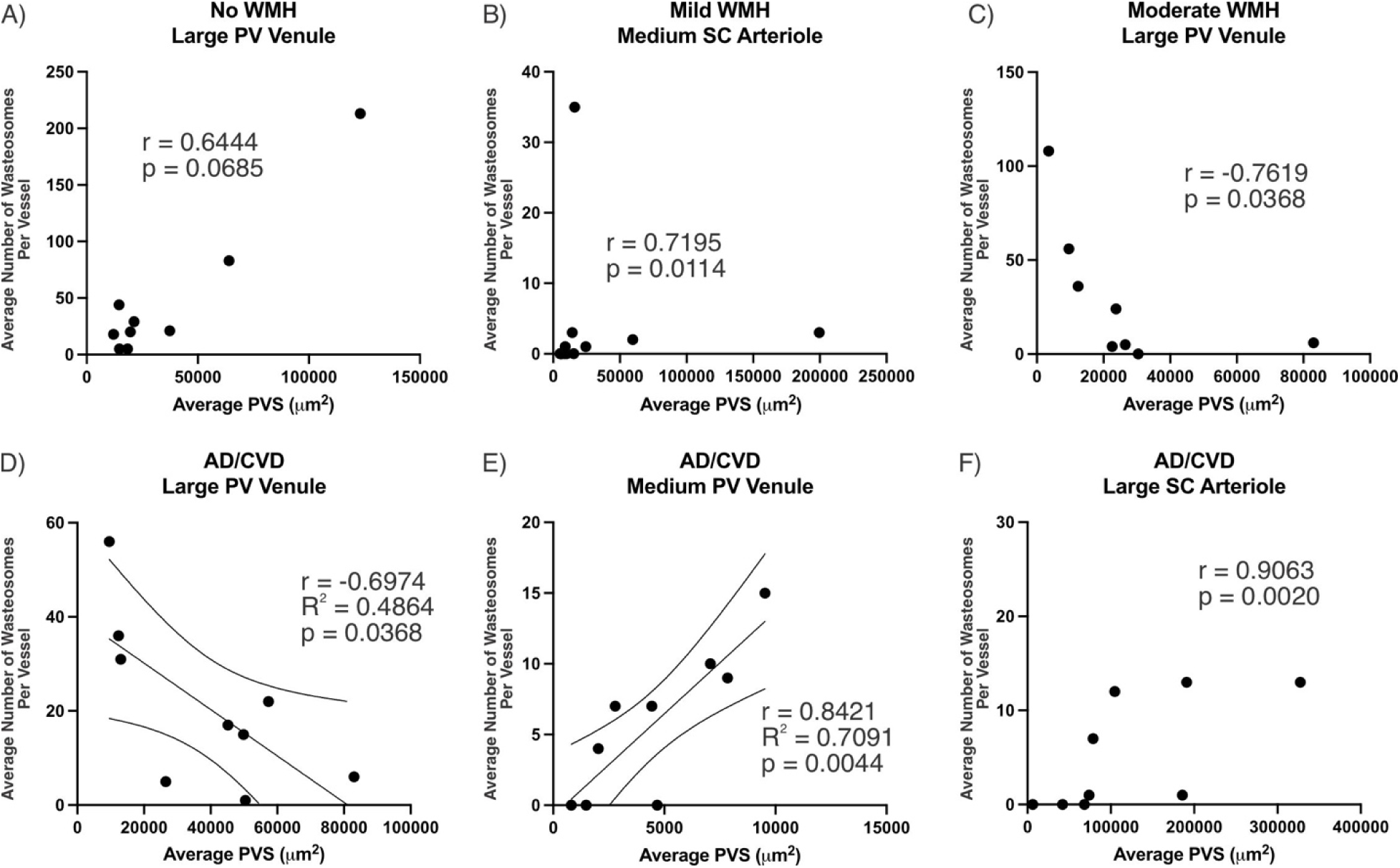
Evidence of a complex relationship between PVS and wasteosome accumulation. A) In specimens with no magnetic resonance imaging (MRI) confirmed white matter hyperintensity (WMH) there was an insignificant positive relationship between perivascular space (PVS) and wasteosome accumulation around large periventricular (PV) venules. B) In specimens with mild WMHs there was a significant positive relationship between PVS and wasteosome accumulation around medium subcortical (SC) arterioles (r = 0.7195, p = 0.0114). C) However, in specimens with moderate WMHs there was a significant negative relationship between PVS and wasteosome accumulation around large PV venules (r = -0.7619, p = 0.0368). D) A negative relationship was also observed in specimens with AD/CVD pathological diagnoses between wasteosome accumulation and PVS of large PV venules (r = -0.6974, R^2^ = 0.4864, p = 0.0368, simple linear regression with 95% confidence interval shown on graph). However, positive relationships between PVS and wasteosome accumulation in specimens with AD/CVD were apparent in E) medium PV venules (r = 0.8421, R2 = 0.7091, p = 0.0044, simple linear regression with 95% confidence interval shown on graph), and F) large SC arterioles (r = 0.9063, p = 0.0020).

## Discussion

Despite first being described in the brain in the 1800’s by Purkinje and Virchow and being well recognized features of the brain’s histological architecture by neuropathologists, wasteosomes remain poorly understood. Originally thought to be of no pathological significance, wasteosomes have largely been thought irrelevant in the study of neurodegenerative disease. However, with advances in technology and the emergence of research into the glymphatic system, wasteosomes are re-emerging in the literature as highly relevant entities in both aging and disease. This study is the first to investigate wasteosome accumulation in the context of PV WMHs in a mixed diagnostic cohort. This study not only better characterizes the localization of wasteosomes in this neuroanatomical region but also raises three key findings warranting further discussion: 1) wasteosome accumulation is increased within PV WMHs; 2) wasteosome accumulation may be independent of WMH scoring but may be linked to disease pathology; and, 3) a dynamic relationship exists between wasteosome accumulation, SI and PVS.

The most significant finding was the preferential accumulation of wasteosomes within PV WMHs relative to NAWM in the combined moderate and PVI specimen groups (Figure 1B,D). The observed increase in wasteosomes may be in response to: 1) an increased production of wasteosomes; 2) a decreased clearance of wasteosomes; or 3) an unknown confound of glymphatic flow causing wasteosomes to preferentially accumulate at neuroanatomical foci where PV WMHs occur. As noted in the introduction, astrocytes are now the cell largely credited for the production of wasteosomes in the central nervous system^2,3,24–33^. While WMHs are often presumed to be of vascular origin and considered features of small vessel disease, the pathology behind WMHs is heterogenous and the etiology cannot be elucidated from imaging alone. Despite this uncertainty, astrocytes are critically implicated in white matter homeostasis, blood brain barrier (BBB) maintenance, and are known to respond to both ischemic and non-ischemic mechanisms of white matter injury^34^. Therefore, it is plausible that the increase in wasteosome accumulation is due to an increase in astrocytic activation and “clean-up” of ischemic or otherwise challenged white matter. Astrocytic wasteosomes have also been postulated to serve as a mechanism for the clearance of dysfunctional mitochondria, further suggesting a connection to metabolic failure and incomplete breakdown of cellular waste^26,28,35^. As noted in the introduction PVI may be considered a finding suggestive of an ischemic or vascular underpinning in PV WMHs. Previously, in these specimens, our group has connected the presence of PVI in PV WMH with microvessel stenosis, enlarged PVSs, increased glial activity and BBB extravasation of fibrinogen^16^. In this study we found no difference between the moderate WMH scored group and the PVI group of specimens (Figure 1C). As noted in Roseborough et al. 2021, lack of PVI does not necessarily imply non-vascular origin^16^. Additionally, in these specimens only PVIs within the same coronal section of the PV WMH were considered, and therefore additional coronal sections may have PVIs and greater underlying vascular pathology^16^. Vascular challenge and subsequent astrocytic activation should be explored as the cause of increased wasteosome accumulation in WMHs in future research.

Vascular pathology may also be involved in the potential reduced clearance of wasteosomes. We hypothesized that increased SI would correlate with increased wasteosome accumulation around blood vessels in a “clogged sink” mechanism of glymphatic drainage. Intriguingly, in medium sized vessels in varying conditions we saw the opposite effect. As noted in the introduction, ISF flow through the glymphatic system is in part driven through the pulsatile nature of arterioles^36,37^. In Figure 3B, we showed an inverse relationship between medium sized PV arteriole SI and wasteosome accumulation in specimens with no pathology diagnosis. It is possible increased SI correlates with decreased pulsatility and therefore fewer wasteosomes shunting towards arterioles with greater SI. Additionally, the “clogged sink” mechanism, may in fact have the opposite effect, by which wasteosomes bypass vessels with greater SI. We were similarly intrigued by the results of the PVS correlations. We hypothesized greater PVS would correlate with greater wasteosome accumulation since those vessels would function as larger glymphatic drains. This relationship potentially exists in large PV venules in specimens with no WMHs (Figure 4A) and medium SC arterioles in specimens with mild WMHs (Figure 4B) but is seemingly reversed to a negative relationship in large PV venules in specimens with moderate WMHs (Figure 4C). Potentially this inverse relationship is occurring as WMH severity is increased; however, future work and a larger sample size is needed to elucidate if this is a consistent finding. Similarly, while this relationship was found in medium PV venules and large SC arterioles in AD/CVD (Figure 4E,F), it was again reversed in large PV venules in AD/CVD. Intriguingly, the mixed AD/CVD was also found to have increased wasteosome accumulation around medium SC arterioles compared to specimens with only a CVD pathology diagnosis (Figure 2O), suggesting a synergistic effect of the comorbid pathology. Previous work has demonstrated increased wasteosomes in both AD^38,39^ and frontotemporal lobar degeneration relative to controls^40^. It is important to note, that our work supports this literature linking wasteosome accumulation to neurodegenerative pathology but also adds to the literature by 1) showing a synergistic accumulation in mixed neurodegenerative and vascular pathology; and 2) further interrogating wasteosome accumulation in relation to specific vessel subtypes. This may also explain why our findings show more modest differences between these groups. It is also important to note that the findings in this study are temporally limited, as all results are depictions of a snapshot in time in fixed tissue, whereas wasteosome accumulation around WMHs and PV blood vessels is likely to be a dynamic and fluid process. Additionally, SI and PVS are only two means of assessing vascular dysregulation in relation to wasteosome accumulation. Other microvessel abnormalities such as lipohyalinosis and cerebral amyloid angiopathy where not assessed and should be considered in future work^41,42^. Additionally, unmatched age between cohorts should be considered as a limitation. The average age at time of death of the AD, CVD, AD/CVD and no pathology diagnosis cohorts were 77.2 years, 72.6 years, 80.4 years and 61.6 years respectively. Moreover, we do not have access to pre-mortem clinical information on these donors to know if they had any comorbidities that may impact glymphatic flow and wasteosome accumulation. Collectively, a complex and dynamic relationship between wasteosome accumulation and different neurodegenerative and neurovascular diseases is only beginning to emerge. Further work with larger sample sizes is needed to validate these relationships in varying diagnostic and WMH scored groups.

Finally, the third option remains possible, that the nature of glymphatic flow independent of the WMHs may have caused wasteosomes to accumulate at the same neuroanatomical location WMHs occurred. However, this leads to the discussion of whether “glymphatic drainage sinks” where wasteosomes accumulate, may exhibit increased susceptibility to the development of WMHs. Studies correlating glymphatic dysfunction and WMHs using the diffusion tensor imaging analysis along the PVS (ALPS)-index are continuously emerging^43–45^. Further, glymphatic system dysregulation has been shown to play a moderator role in the relationship between WMHs and cognitive impairment^46^. A separate study found that PV WMHs were mainly attributed to damage to the venous system due to glymphatic dysfunction, whereas deep WMHs could occur as a result of ischemic injury and glymphatic dysfunction^47^. Our results demonstrating wasteosomes preferentially accumulate around large PV venules (Figure 1E,F) further suggests a key relationship between the wasteosome system and PV venous drainage.

Further, by normalizing these findings to PVS, we showcase that this relationship is considerably complex and not merely a function of differing PVS area (Figure 1G,H). Collectively, there appears to be a dynamic relationship between WMHs and glymphatic dysfunction. It remains possible that glymphatic dysregulation results in wasteosome accumulation which precipitates or exacerbates WMH pathology. Future work should continue to characterize wasteosomes in white matter lesions in a variety of neurodegenerative, neurovascular and mixed pathologies.

## Conclusions

This study is the first to investigate wasteosome accumulation in PV WMHs in a mixed diagnostic cohort. The results from this study better characterize the localization of wasteosomes in PV white matter and demonstrate a relationship in which wasteosomes accumulate in PV WMHs, are influenced by disease pathology and exhibit a dynamic relationship with vessel subtype, SI and PVS. Wasteosomes represent a novel means of investigating glymphatic dysregulation in post-mortem tissue and wasteosome accumulation in neurodegenerative pathology is emerging as an important biomarker of disease pathology. Further work is needed to understand the mechanisms and implications of wasteosome accumulation and glymphatic dysregulation.

